# Shape Recovery of Deformed Biomolecular Droplets: Dependence on Condensate Viscoelasticity

**DOI:** 10.1101/2021.01.20.427476

**Authors:** Huan-Xiang Zhou

## Abstract

A theoretical study on the shape dynamics of phase-separated biomolecular droplets is presented, highlighting the importance of condensate viscoelasticity. Previous studies on shape dynamics have modeled biomolecular condensates as purely viscous, but recent data have shown them to be viscoelastic. Here we present an exact analytical solution for the shape recovery dynamics of deformed biomolecular droplets. The shape recovery of viscous droplets has an exponential time dependence, with the time constant given by the “viscocapillary” ratio, i.e., viscosity over interfacial tension. In contrast, the shape recovery dynamics of viscoelastic droplets is multi-exponential, with shear relaxation yielding additional time constants. During shape recovery, viscoelastic droplets exhibit shear thickening (increase in apparent viscosity) at fast shear relaxation rates but shear thinning (decrease in apparent viscosity) at slow shear relaxation rates. These results highlight the importance of viscoelasticity and expand our understanding of how material properties affect condensate dynamics in general, including aging.

## INTRODUCTION

Phase-separated biomolecular condensates exhibit different extents of liquidity, which may be crucial for cellular functions and correlated with diseases such as neurodegeneration and cancer (Alberti and Dormann, 2019; Alberti et al., 2019; Hubstenberger et al., 2013; Li et al., 2020; Roden and Gladfelter, 2020; Yamasaki et al., 2020; Yu et al., 2020). Driven by interfacial tension (or capillarity), condensates that are more liquid-like retain a spherical shape and hence appear as micro-sized droplets. The material properties of many condensates also evolve over time and become more solid-like (sometimes referred to as “aging”) (Feng et al., 2019; Gui et al., 2019; Jawerth et al., 2020; Patel et al., 2015; Woodruff et al., 2018; Zhang, 2020). A simple indication of liquidity is the tendency of deformed droplets to recover their spherical shape. In all the studies so far on the dynamics of such shape changes, biomolecular condensates have been modeled as purely viscous, reporting viscosities that are orders of magnitude higher than that of water (Brangwynne et al., 2009; Brangwynne et al., 2011; Elbaum-Garfinkle et al., 2015; Feric et al., 2016; Hubstenberger *et al.*, 2013; Patel *et al.*, 2015; Zhang et al., 2015). However, recent work has shown that condensates are viscoelastic (Alshareedah et al., 2021; Ghosh et al., 2021; Jawerth *et al.*, 2020; Jawerth et al., 2018; Zhou, 2020; 2021). In viscoelastic fluids, shear relaxation is not instantaneous (Zhou, 2021) and its effects, when coupled with viscocapillary effects, can lead to unexpected observations. In this work we present a theoretical study on the effects of viscoelasticity on the shape recovery dynamics of deformed biomolecular droplets.

Nearly all previous studies on the shape dynamics of biomolecular condensates have dealt with the fusion of droplets when they come into contact (Alshareedah et al., 2019; Boeynaems et al., 2019; Brangwynne *et al.*, 2009; Brangwynne *et al.*, 2011; Caragine et al., 2018; Elbaum-Garfinkle *et al.*, 2015; Feric *et al.*, 2016; Ghosh and Zhou, 2020; Gui *et al.*, 2019; Hubstenberger *et al.*, 2013; Patel *et al.*, 2015; Wang et al., 2018; Zhang *et al.*, 2015). However, in one study, Hubstenberger et al. (Hubstenberger *et al.*, 2013) reported observations on both droplet fusion and shape recovery of grP-bodies formed in the cytoplasm of *Caenorhabditis elegans* oocytes. Shape recovery after mechanical stress-induced elongation was faster than droplet fusion by two orders of magnitude, which was suggested as indicating elasticity of these ribonucleoprotein condensates. While theoretical results of viscous fluids have been used for qualitative as well as quantitative analysis of droplet fusion dynamics (Brangwynne *et al.*, 2009; Brangwynne *et al.*, 2011; Elbaum-Garfinkle *et al.*, 2015; Feric *et al.*, 2016; Ghosh and Zhou, 2020; Hubstenberger *et al.*, 2013; Zhang *et al.*, 2015) and a theoretical model of viscoelastic fluids has been introduced to treat deformation of droplets under external force (Jawerth *et al.*, 2018; Zhou, 2020), up to now no theory has been presented for shape recovery of deformed biomolecular droplets.

However, the problem of shape recovery has long received attention in the fluid-dynamics literature. Prosperetti (Prosperetti, 1977; Prosperetti, 1980) presented a full analytical solution to the recovery dynamics of viscous droplets from a small-amplitude deformation. Phenomenological models have also been introduced to propose theoretical results for shape recovery, also known as deformation retraction, of viscoelastic droplets (Minale, 2010; Yu et al., 2004). This problem has also been tackled by numerically solving the fluid-dynamics equations (Hooper et al., 2001; Mukherjee and Sarkar, 2010; Verhulst et al., 2009b). There have also been a number of experimental studies on the deformation retraction of polymer blends (Ali and Prabhu, 2019; Luciani et al., 1997; Tretheway and Leal, 2001; Verhulst et al., 2009a), often indicating the effects of viscoelasticity.

The high viscosities reported in *in vitro* studies of biomolecular condensates (relative to water) translate into a significant viscosity ratio, *η*_II_/*η*_I_, between the dense droplet phase (indicated by index II; Fig. 1) and the exterior dilute bulk phase (indicated by index I). On the other hand, Caragine et al. (Caragine *et al.*, 2018) suggested a case with *η*_II_/*η*_I_ ≪ 1. They noted the even higher viscosity of the nucleoplasm surrounding nucleoli, a type of biomolecular condensates specializing in ribosomal biogenesis, than that of droplets formed by a main nucleolar protein (Feric *et al.*, 2016). In addition, polymer blends may phase separate to produce dilute-phase “bubbles” in dense coacervates (Ali and Prabhu, 2019), also leading to *η*_II_/*η*_I_ ≪ 1. The viscosity ratio is of theoretical interest because in the limiting cases of *η*_II_/*η*_I_ = ∞ and 0 one can neglect the fluid dynamics in either the exterior region (i.e., “I”) or interior region (i.e., “II”), significantly simplifying the mathematics. The viscosity ratio may also affect how viscoelasticity affects shape dynamics.

**FIG. 1.**
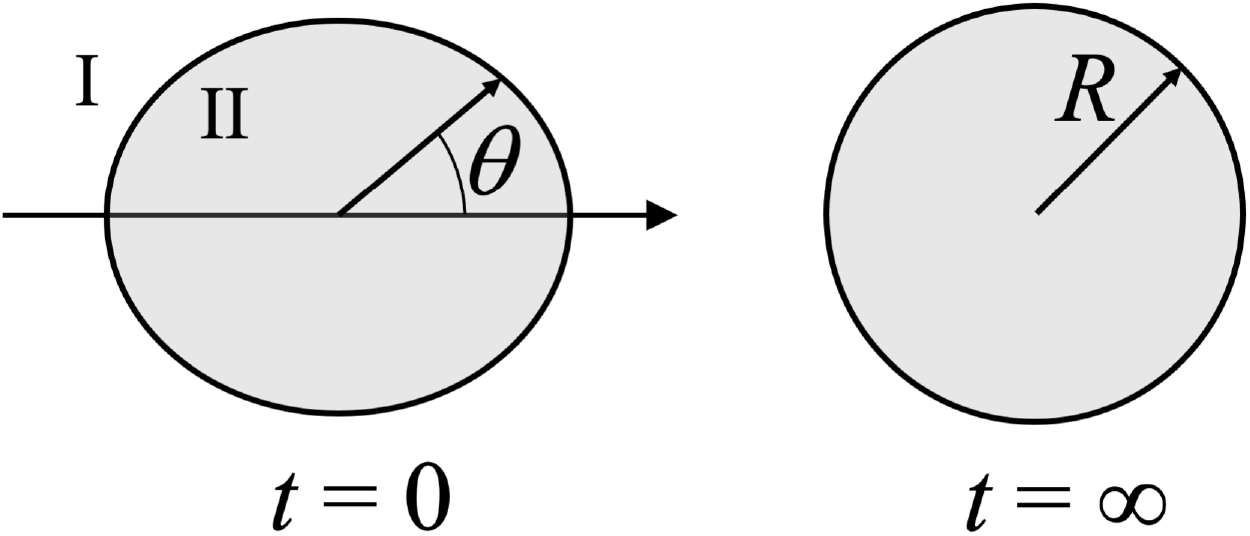
Illustration of the shape recovery of a deformed droplet. Left: initial deformation; right: fully recovered spherical shape.

Here we develop an analytical solution for the shape recovery dynamics of deformed viscoelastic droplets. First we summarize the solution of Prosperetti for viscous fluids (Prosperetti, 1977; Prosperetti, 1980) and develop a new solution that is amenable to generalization to viscoelastic fluids. The solution for viscoelastic fluids is then presented. Readers not interested in technical details can skim over these two sections and focus on the next section, where we report shape recovery curves for selected values of viscosities and shear relaxation rates, to illustrate rich effects of viscoelasticity. In particular, during shape recovery, viscoelastic droplets exhibit shear thickening (increase in apparent viscosity) at fast shear relaxation rates but shear thinning (decrease in apparent viscosity) at slow shear relaxation rates. Moreover, our analytical results show that shear relaxation is a new rate-limiting mechanism for shape dynamics, although experimental verification can be challenging. The paper ends with concluding remarks. We also place additional technical details in four Appendices.

## INTERFACE SHAPE RCOVERY: NEWTONIAN FLUIDS

Our main interest is biomolecular droplets formed by phase separation. Due to vast differences in macromolecular concentrations between the droplet phase and the surrounding bulk phase, the viscosity in the droplet phase is much higher than the counterpart in the bulk phase, i.e., *η*_II_ ≫ *η*_I_. Neglecting the viscosity of the bulk phase (i.e., approximating the latter as an ideal fluid) leads to significant simplification of the solution of the fluid-dynamics problem. We then denote the viscosity of the droplet phase as *η*, without the subscript “II”.

For some polymer blends, phase separation leads to polymer-poor “bubbles” forming in the coacervate matrix (Ali and Prabhu, 2019). In this case, *η*_I_ ≫ *η*_II_; we then neglect *η*_II_ and denote the viscosity of the coacervate phase as *η*, without the subscript “I”. We only retain the subscripts I and II when considering the full problem where both interior and exterior fluid dynamics are treated.

### Fluid-dynamics problem

The shape change of a liquid droplet inside an ideal-fluid medium is dictated by the interior fluid dynamics, which we model by the (generalized) Navier-Stokes equations. The first of these equations expresses mass conservation, which for an incompressible fluid has the form

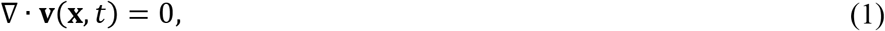

where **v**(**x**, *t*) is the fluid velocity at position **x** and time *t*, and **∇** denotes the gradient operator. The second of the Navier-Stokes equations expresses momentum conservation,

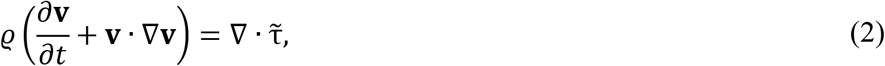

where *ϱ* is the fluid density and 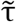 is the stress tensor. Closure of this equation requires a constitutive relation for 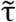. One contribution to 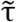 is hydrostatic pressure (*p*); the remaining contribution comes from fluid viscosity:

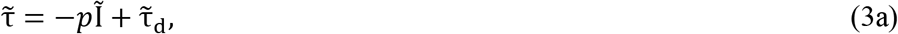

where Ĩ is the unit tensor. For purely viscous, or Newtonian fluids, the second contribution is given by

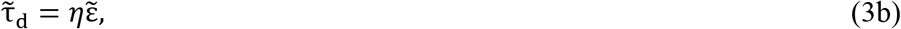

where *η* is the fluid viscosity, and

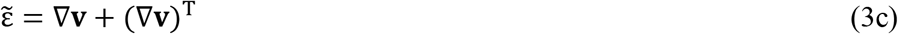

is the symmetrized shear-rate tensor; the superscript “T” denotes transpose. The momentum equation then becomes

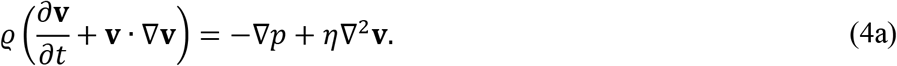

Biomolecular condensates have very high viscosity. Then the inertial terms on the left-hand-side of Eq. (4a) can be neglected, leading to

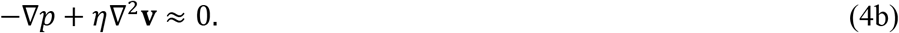

The reduction of Eq. (4a) to Eq. (4b) is similar to the reduction, at high frictions, of the equation of motion for a Langevin particle to one representing Brownian dynamics.

To solve Eqs. (1) and (4a) [or (4b)], we have to specify boundary conditions. One is a “kinematic” boundary condition, which expresses the fact that fluid motions lead to changes in the shape of the interface between the droplet and bulk phases. If the interface at time *t* is specified by the condition

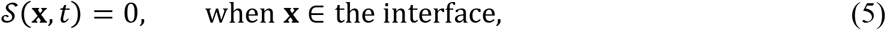

then the kinematic boundary condition is

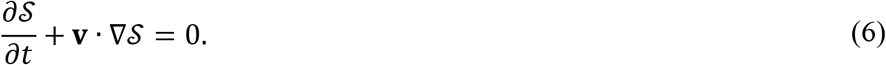

The outward unit normal vector of the interface is given by

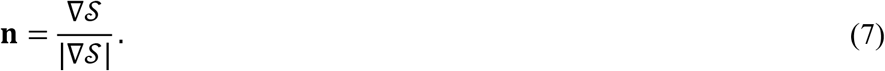

The remaining boundary conditions express force balance at the interface:

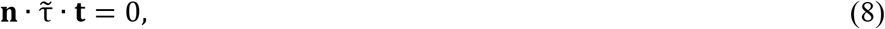

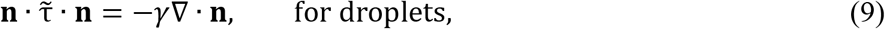

where **t** denotes a unit vector along a tangential direction of the interface, and *γ* is the interfacial tension. Note that, due to the much higher viscosity inside biomolecular droplets than in the surrounding bulk phase, the effect of the velocity field in the bulk phase has been neglected. A constant pressure in the bulk phase can be accounted for by subtracting it from the interior pressure. Therefore, the two force-balance boundary conditions involve only the interior stress tensor.

In the case of ideal-fluid bubbles in a Newtonian-fluid medium, the shape change is dictated by the exterior fluid dynamics. All but one of the above equations still hold when the symbols denote exterior properties. The one exception is a sign change in the boundary condition of Eq. (9):

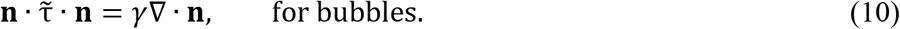

The problem at hand is to find the interface shape function 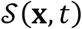 at any time *t*, given the initial shape function 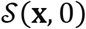 (and its initial rate 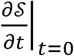 if necessary). Due to the interfacial tension, a deformed droplet will recover its spherical shape; we refer to this process shape recovery. Other terms used in the literature include deformation retraction or relaxation.

### Solution of Prosperetti

Prosperetti (Prosperetti, 1977) solved the shape recovery problem for small-amplitude deformation. That is, the initial shape of the interface bewteen a droplet and the bulk phase (approximated as ideal fluid) was assumed to be a small deformation from a sphere with radius *R*. Using spherical coordinates (*r, θ, ϕ*) for the position **x** (with origin at the center of the reference sphere), the radial distance of the interface at polar angle *θ* and azimuthal angle *ϕ* and at time *t* can be written as (Fig. 1)

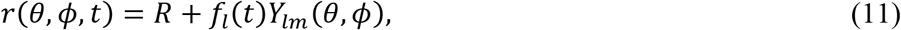

where *Y_lm_*(*θ, ϕ*) is a spherical harmonic. Because of the orthogonality of sphercial harmonics, we can later add up the results for individual spherical harmonics to obtain a complete solution. The amplitude *f_l_*(*t*) satisfies the following integro-differential equation:

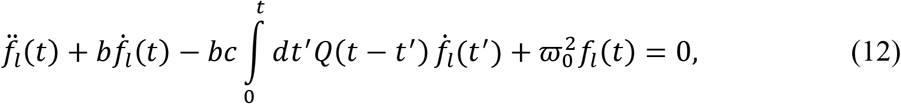

where dots denote differentiation with respect to time,

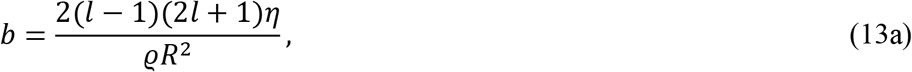

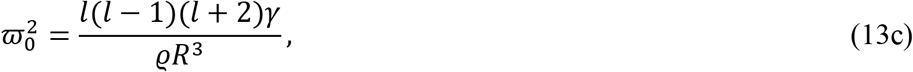

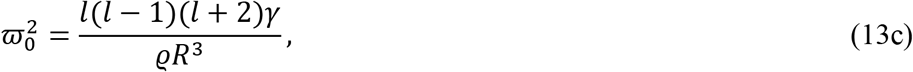

and for the function *Q*(*t*) only its Laplace transform is known. Denoting the Laplace transform of any function of time, *g*(*t*), as *ĝ*(*s*), we have

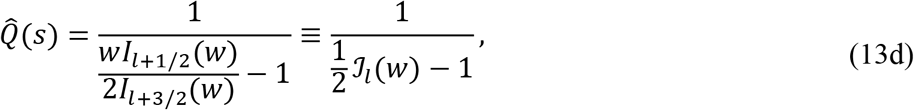

where 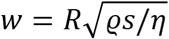 and *I*_*l*+1/2_(*x*) are modified Bessel functions of the first kind.

Prosperetti (Prosperetti, 1977) considered the long-time asymptotic behavior of *f_l_*(*t*), where *f_l_*(*t*) ≈ *C*(*t*)*e*^−*ν_l_t*^ with *C*(*t*) becoming a finite constant as *t* → ∞, leading to an equation for *ν_l_* that is identical to the classical result of Chandrasekhar (Chandrasekhar, 1961) and Reid (Reid, 1960) using normal-mode analysis. This analysis has now been extended to viscoelastic droplets in an ideal fluid (Brenn and Teichtmeister, 2013; Khismatullin and Nadim, 2001).

Here our concern is the initial value problem, i.e., the time dependence of *f_l_*(*t*) starting from *t* = 0. We are particularly interested in the high-friction regime. In this regime, we neglect the second derivative, 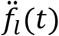, in Eq. (12), similar to reducing the Langevin equation to one representing Brownian dynamics. Furthermore, high friction means *w* → 0 and hence we approximate 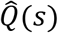 as

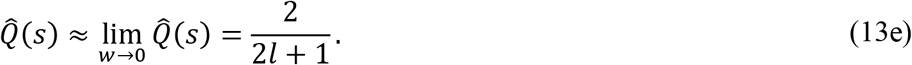

This result is independent of *s*, meaning that *Q*(*t*) is a delta function of time. The solution of Eq. (12) is an exponential function,

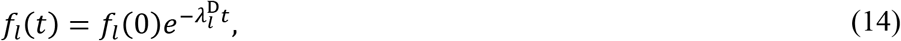

with the recovering rate given by

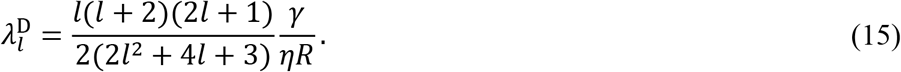

For ideal-fluid bubbles, Eq. (12) still holds, but with

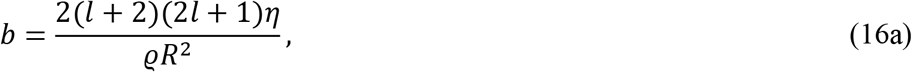

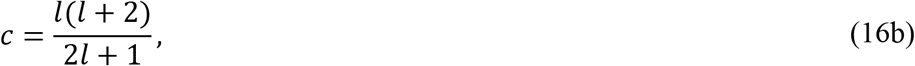

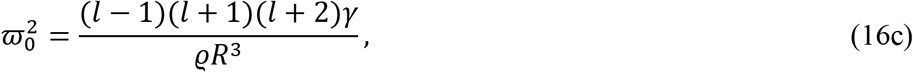

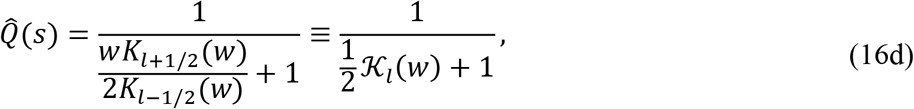

where *K*_*l*+1/2_(*x*) are modified Bessel functions of the second kind. Again our interest is the high-friction regime, where *w* → 0 and 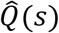 of Eq. (16d) approaches the same limit as given by Eq. (13e). The solution for *f_l_*(*t*) remains an exponential function, with the recovery rate now given by

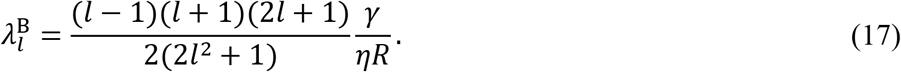

Prosperetti (Prosperetti, 1980) also solved the full problem where both interior and exterior fluid dynamcis are treated, i.e., without invoking *η*_II_ » *η*_I_ or *η*_II_ « *η*_I_. Equation (12) becomes

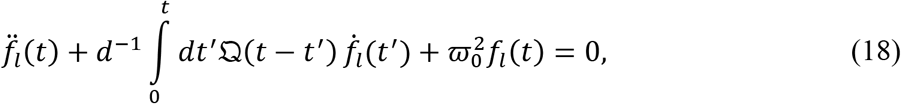

where

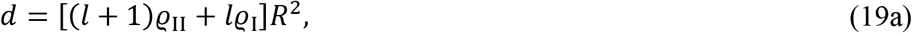

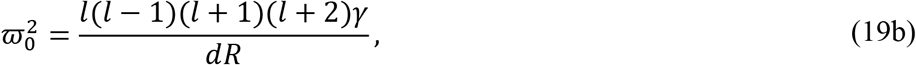

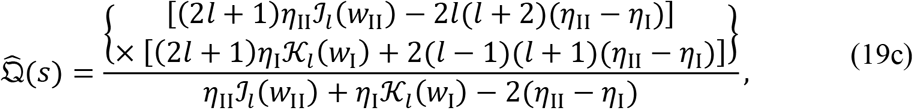

with subscripts “II” and “I” denoting quanties in the interior and exterior of the droplet. In the high-friction regime,

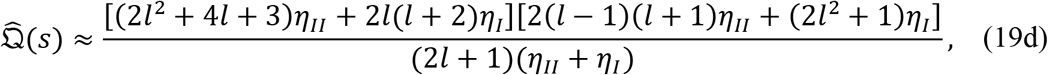

and *f_l_*(*t*) is an exponential funciton of time, with the recovery rate given by

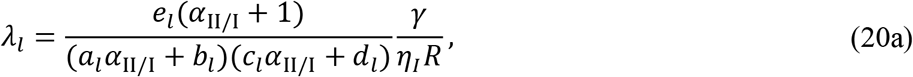

where

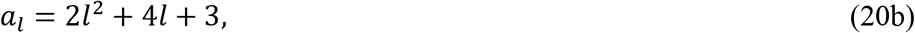

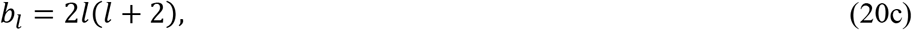

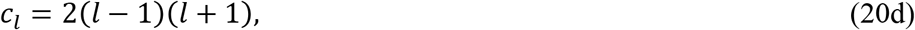

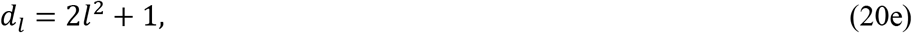

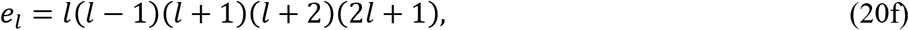

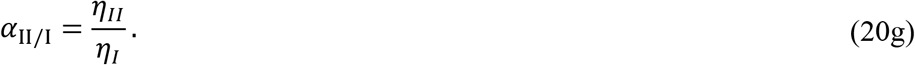

One can easily verify that *λ_l_* reduces to 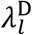 when *α*_II/I_ ≫ 1 and to 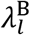 when *α*_II/I_ ≪ 1.

### New solution

Here present a new solution for the shape recovery dynamics in the high-friction regime. This solution easily lends itself to generalization from viscous to viscoelastic fluids. The derivation here for droplets largely follows the solution of a related fluid-dynamics problem (Zhou, 2020). We assume that the shape deformation is axisymmetric (Fig. 1). Equation (11) then reduces to

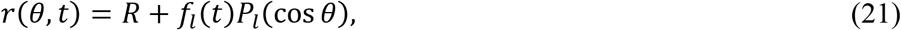

where *P_l_*(*x*) is a Legendre polynomial. Note that the lowest *l* should be 2, since *l* = 0 corresponds to a change in droplet radius and thus a violation of volume conservation demanded by fluid incompressibility, and *l* = 1 corresponds to a translational motion of the entire interface. We will keep terms only up to the first order in *f_l_*(*t*); to this order, the volume of the droplet is a constant when *f_l_*(*t*) changes over time.

The interface shape function can be identified as

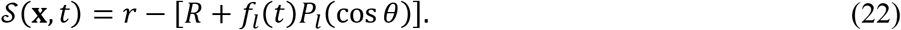

The outward normal vector of the interface [Eq. (7)] can be found as

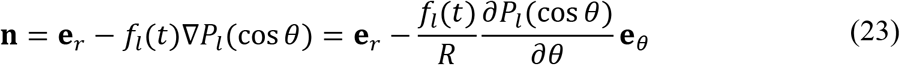

to the first order in *f_l_*(*t*). The divergence of **n** is

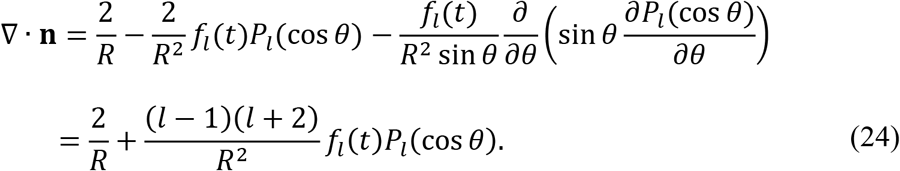

The fluid velocity **v**(**x**, *t*) is of the same order as *f_l_*(*t*). Similarly, the pressure should only deviate from the static pressure that balances the interfacial tension of a spherical droplet by an amount, *δp*(**x**, *t*), that is of the same order as *f_l_*(*t*):

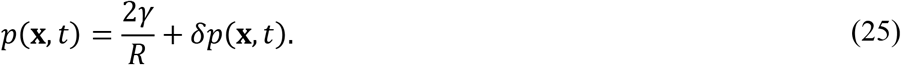

The kinematic boundary condition [Eq. (6)] yields

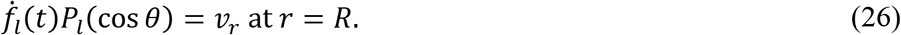

Subscripts *r* and *θ* denote the components of the velocity. The force-balance boundary conditions [Eqs. (8) and (9)] lead to

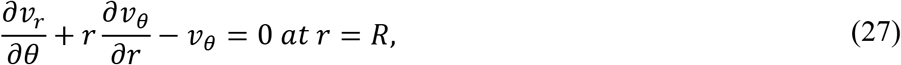

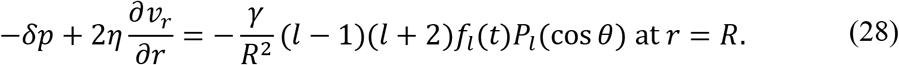

The velocity and pressure fields for an axisymmetric problem can be expressed in terms of the stream function *Ψ*(*r, θ, t*). For the interior problem considered here for droplets, the stream function has the form (Leal, 2007)

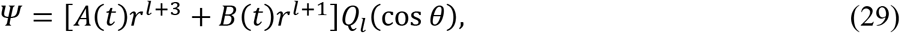

where *A*(*t*) and *B*(*t*) are coefficients to be determined by the boundary conditions, and *Q_l_*(*x*) are related to Legendre polynomials *P_l_*(*x*) via

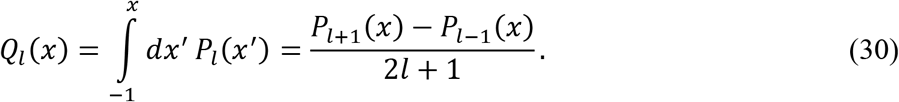

The components of the velocity and the pressure are given by

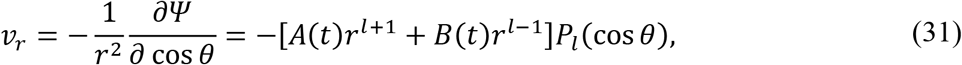

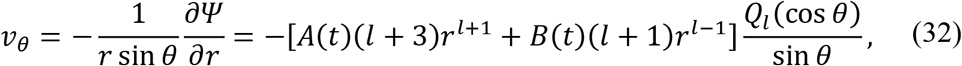

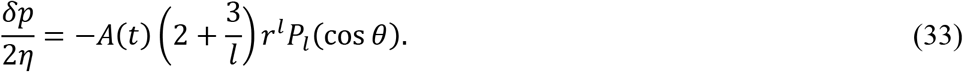

Substituting Eqs. (31) and (32) into Eq. (27) leads to

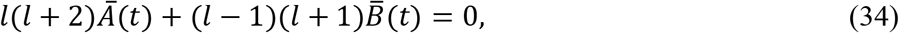

where *Ā*(*t*) = *A*(*t*)*R^1^* and 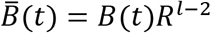. Similarly, using Eqs. (31) and (33) in Eq. (28) leads to

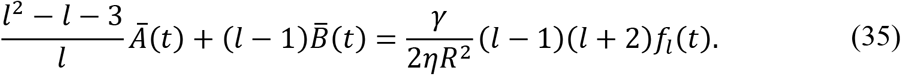

The results for *Ā*(*t*) and 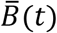 are:

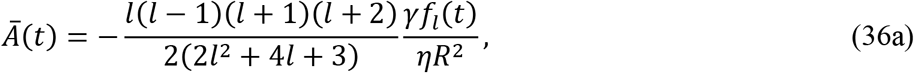

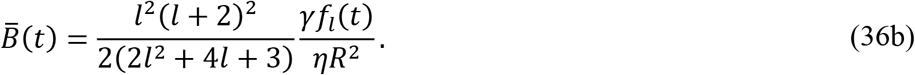

When these last results are inserted into Eq. (31), we find

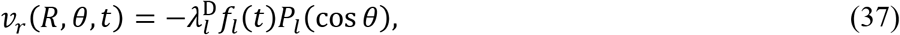

where 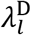 is given by Eq. (15). Substituting Eq. (37) in Eq. (26) yields

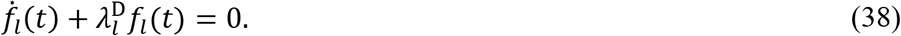

The solution of this last equation is the exponential function in Eq. (14).

In preparation for the generalization to viscoelastic fluids, let us solve the problem again, now using Laplace transforms. The governing equations (1) and (4b) and the forcebalance boundary conditions [Eqs. (27) and (28)] have the same form in Laplace space as they have in the time domain, as does the general form of the stream function [Eq. (29)]. Therefore Eqs. (34) and (35) take the same form in Laplace space:

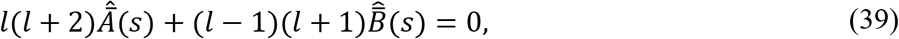

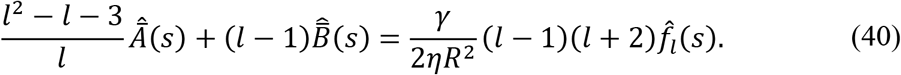

The kinematic boundary condition [Eq. (26)] now takes the form

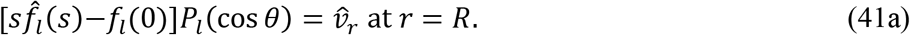

Using Eq. (31) in the preceding equation, we find

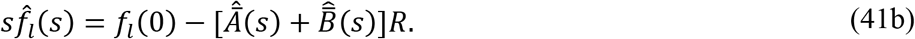

Solving Eqs. (39), (40), and (41b), we find

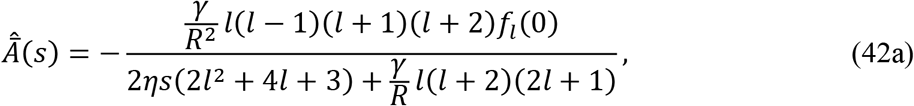

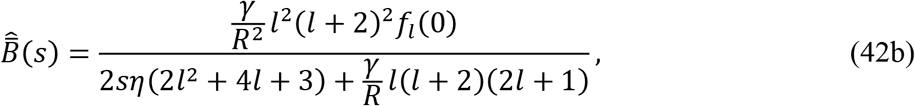

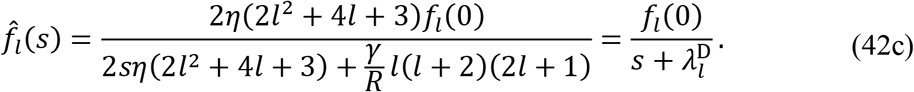

The inverse Laplace transform of the foregoing 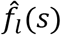 is given by Eq. (14).

For the exterior problem appropriate for bubbles, the stream function is (Leal, 2007)

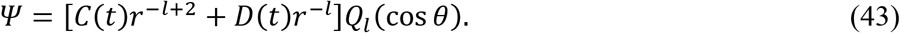

Analogous to Eqs. (31) – (33), we find the velocity and pressure to be

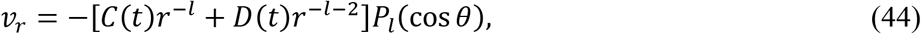

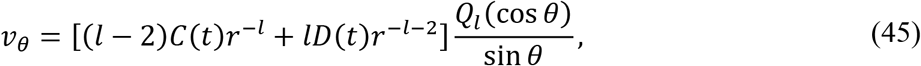

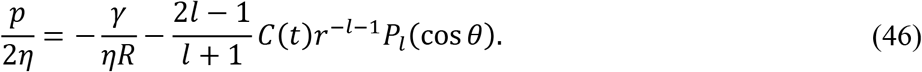

The force-balance boundary conditions [Eqs. (8) and (10), appropriate for bubbles] lead to

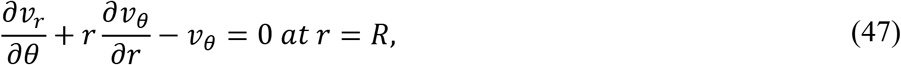

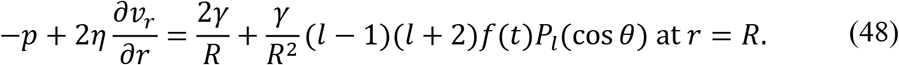

Substituting Eqs. (44) – (46) into Eqs. (47) and (48) and expressing the results in Laplace space, we find

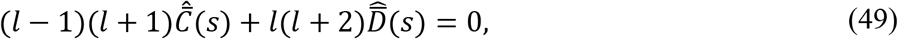

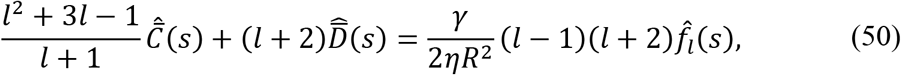

where 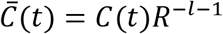 and 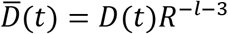. Using Eq. (44) in Eq. (41b), the kinematic boundary condition in Laplace space, we obtain

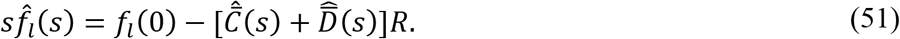

Solving the last three equations, we find

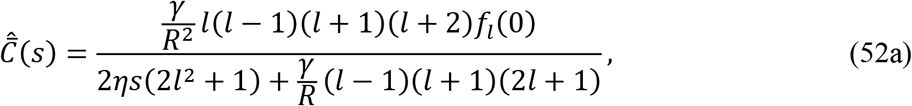

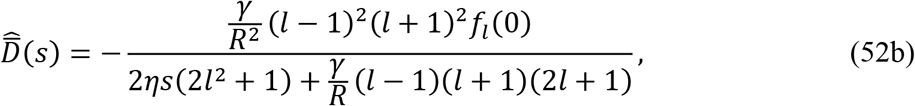

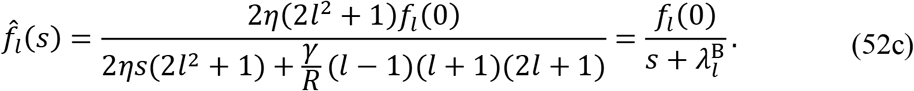

This 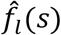 corresponds an exponential function of time, with the recovery rate 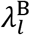 given by Eq. (17).

We also solve the full problem where both interior and exterior fluid dynamcis are treated. The derivation is presented in Appendix A, and the final reuslt for the shape deformation is

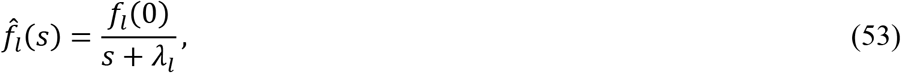

where the recovery rate *λ_l_* is given by Eq. (20a).

## INTERFACE SHAPE RCOVERY: VISCOELASTIC FLUIDS

### Constitutive relation for viscoelastic fluids

In Newtonian fluids, shear relaxation occurs instantaneously, and hence the stress responds only to the strain rate at the same moment [see Eq. (3b)]. For later reference, we present this constitutive relation in Laplace space:

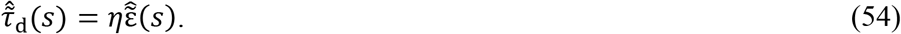

In biomolecular condensates and other complex materials, shear relaxation occurs at a finite rate (Ali and Prabhu, 2019; Jawerth *et al.*, 2020; Verhulst *et al.*, 2009a; Zhou, 2020; 2021), and consequently the stress depends on the entire history of the strain rate. Limiting to small strain rates so that the relation between 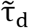 and 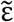 remains linear, Eq. (3b) is generalized to (Zhou, 2021)

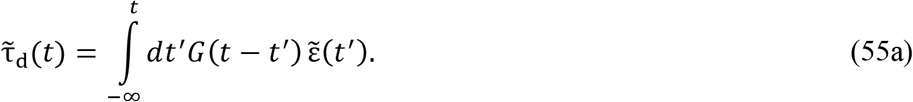

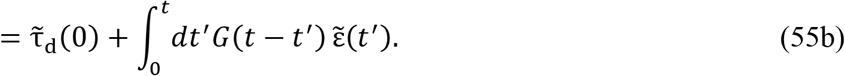

The function *G*(*t*) introduced above is the shear relaxation modulus. For our problem at hand, the strain rate starts at *t* = 0 [i.e., 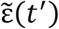 = 0 for *t*’ < 0], and hence

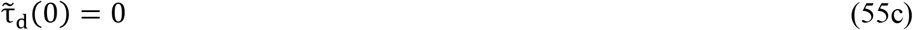

and Eq. (55b) becomes

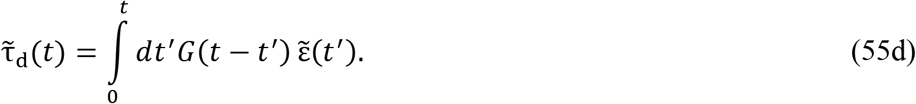

Note that we must also have *G*(*t′*) = 0 for *t′* < 0 since otherwise Eq. (55a) would mean that future strain rates [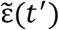 at *t′* > *t*] would affect the present stress 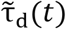. Taking the Laplace transform of Eq. (55d), we obtain a simple constitutive relation between 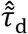 and 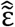:

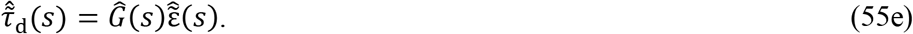

In comparison to Eq. (54) for Newtonian fluids, we see that the only difference is that *η* is now replaced with *Ĝ*(*s*). Therefore, in Laplace space, the Navier-Stokes equations can be generalized to viscoelastic fluids by simply replacing *η* with *Ĝ*(*s*)! Correspondingly, the solution for Newtonian fluids can be transformed to the solution for viscoelastic fluids by the same replacement. We will present this solution in the next subsections.

The Fourier transform of *G*(*t*) defines the complex shear modulus, *G*\*(*ω*), where *ω* is the angular frequency. We have the following relation:

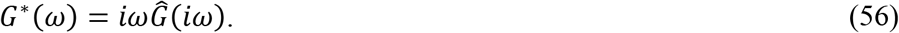

where *i* is the unit imaginary number.

We now introduce various models of linear viscoelasticity. The constitutive relation, Eq. (3b), for Newtonian fluids corresponds to a delta function for the shear relaxation modulus:

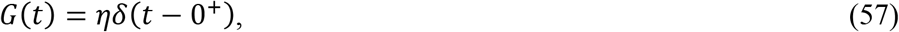

where 0^+^ indicates a peak position that is beyond *t* = 0 by an arbitrarily small positive amount. The Laplace transform of this *G*(*t*) is *η*, thus validating that Eq. (55e) properly reduces to Eq. (54) when the fluids are Newtonian. In the so-called Maxwell model, shear relaxation is an exponential function of time, with a time constant *τ*^sr^,

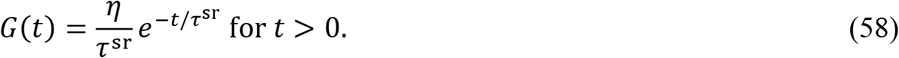

The Laplace transform is

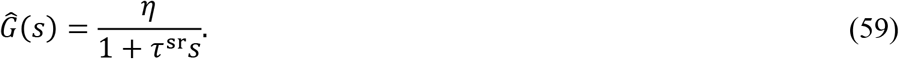

In the limit *τ*^sr^ → 0, the Maxwell model reduces to a Newtonian fluid, consistent with the latter’s instantaneous shear relaxation. A linear combination of a Newtonian fluid and the Maxwell model,

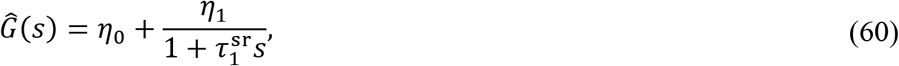

is the Jeffreys model, which can also be seen as the linearized version of the Oldroyd B model. When the Newtonian component of Eq. (60) is also generalized to be Maxwellian, i.e.,

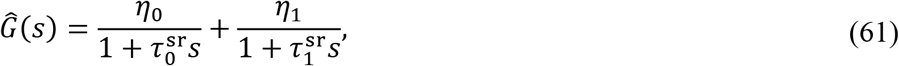

we obtain the Burgers model. In Eq. (61), we assume 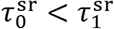. The value of *Ĝ*(*s*) at *s* = 0 will be particularly useful. We call it the zero-frequency viscosity and denote it as

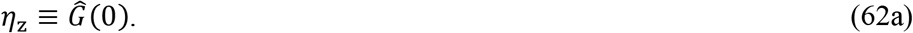

For the Jeffreys and Burgers models, the zero-frequency viscosity is

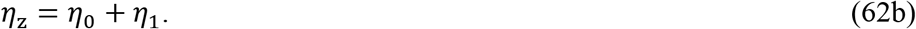

We can represent an arbitrary shear relaxation modulus by adding more and more Maxwellian components. However, these more complicated models rarely are useful.

### Shape recovery dynamics: viscoelastic droplets in an ideal fluid

By substituting the shear relaxation modulus *Ĝ*(*s*) for the viscosity *η* in Eq. (42c), we obtain the shape deformation amplitude, 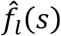, for viscoelastic droplets,

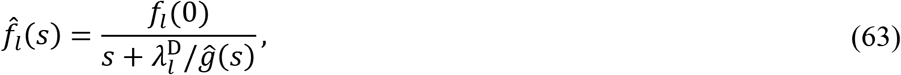

where 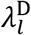 is given by Eq. (15) but with *η* now replaced by *η_z_* = *Ĝ*(0) [eq (62a)], i.e.,

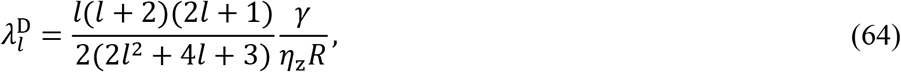

and

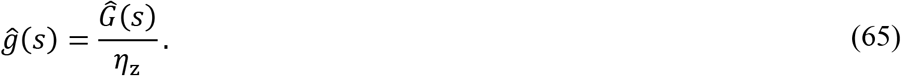

By definition, *ĝ*(0) = 1. For Newtonian fluids, *ĝ*(*s*) = 1 for all *s*. Note that the area under the *f_l_*(*t*) vs. *t* curve, given by 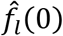, is

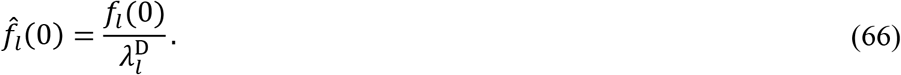

With this result, we can interpret 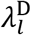 as the mean recovery rate, in that the exponential function 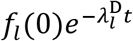 has the same area as *f_l_*(*t*). Equation (66) is derived from Eq. (63) using *ĝ*(0) = 1. Because Eq. (66) holds irrespective of *ĝ*(*s*), the area of the *f_l_*(*t*) curve must be conserved when the parameters, in particular the time constant of shear relaxation, in *ĝ*(*s*) or even the functional forms of *ĝ*(*s*) are varied.

### Shape recovery dynamics: ideal-fluid bubbles in a viscoelastic medium

Similarly, by substituting the shear relaxation modulus *Ĝ*(*s*) for the viscosity *η* in Eq. (52c), we obtain the shape deformation amplitude, 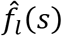, for ideal-fluid bubbles in a viscoelastic medium,

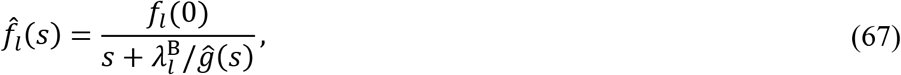

where 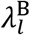 is given by Eq. (17) but with *η* now replaced by *η_z_*,

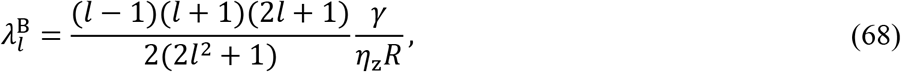

and represents the mean recovery rate that corresponds to the area under the *f_l_*(*t*) vs. *t* curve.

### Shape recovery dynamics: viscoelastic fluids

For the shape recovery of a deformed viscoelastic droplet in a viscous fluid, by substituting *Ĝ*_II_(*s*) for the interior viscosity *η*_II_ in *λ_l_* that is given by Eq. (20a) and appears in Eq. (53), we obtain

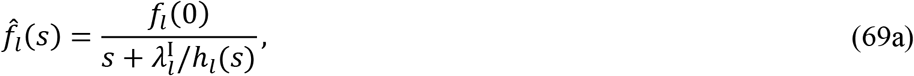

where

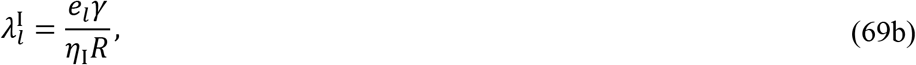

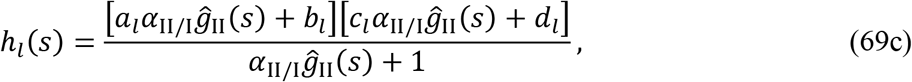

with *a_l_, b_l_, c_l_, d_l_*, and *e_l_* given by Eqs. (20b–f). Furthermore, *α*_II/I_ is the viscosity ratio given by Eq. (20g) but *η*_II_ now represents the zero-frequency viscosity of the viscoelastic droplet, and

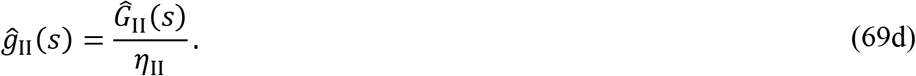

At *s* = 0, we find

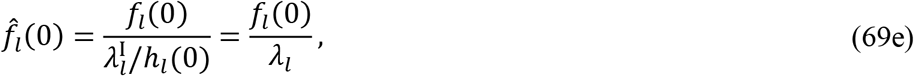

where *λ_l_* is given by Eq. (20a). Equation (69e) means that the area under the *f_l_*(*t*) curve is conserved for any *ĝ*_II_(*s*), including the one, *ĝ*_II_(*s*) = 1, for Newtonian fluids.

The foregoing results can also apply to the reverse case, i.e., a viscous droplet in a viscoelastic medium, after the following changes: replacing 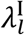 with 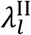 and *ĝ*_II_(*s*) with *ĝ*_I_(*s*), and swapping *a_l_* with *b_l_* and *c_l_* with *d_l_* in Eq. (69c). Finally the shape recovery dynamics of the most general case, i.e., a viscoelastic droplet in a viscoelastic medium, can be obtained by further modifying *λ_l_* in Eq. (53). This time, the modification entails substituting both *Ĝ*_II_(*s*) for the interior viscosity *η*_II_ and *Ĝ*_I_(*s*) for the exterior viscosity *η*_I_ in the expression of *λ_l_* given by Eq. (20a).

## ILLUSTRATIVE RESULTS AND DISCUSSION

The analytical solution in the preceding two sections predicts rich dynamic behaviors of deformed droplets during their shape recovery. Here we present illustrative results, to motivate experimental studies into the effects of viscoelasticity on shape dynamics of biomolecular condensates, and demonstrate the potential importance of the present analytical solution in fitting experimental data and in validating numerical solutions.

### Comparison between droplet fusion and shape recovery

It is interesting to compare the dynamics of the two kinds of shape changes: fusion of two droplets and recovery of a deformed droplet. The half-length, *L*_fus_(*t*), of two fusing viscous droplets, initially with equal radius *R*, in an ideal-fluid medium is approximately given by (Ghosh and Zhou, 2020)

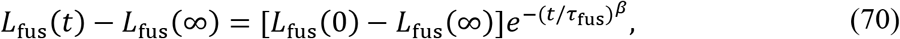

where *β* = 1.5 and

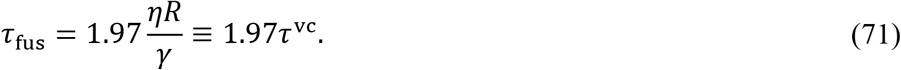

The last identity defines the viscocapillary time *τ^vc^*. For the shape recovery of a deformed droplet, the half-length is given by [Eq. (14)]

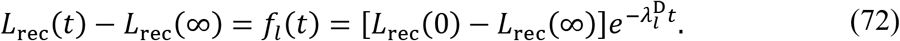

The relaxation rate for *l* = 2 and 4 are [Eq. (15)]

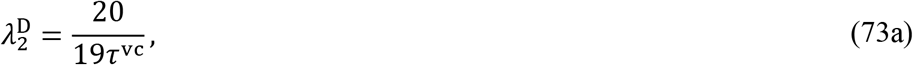

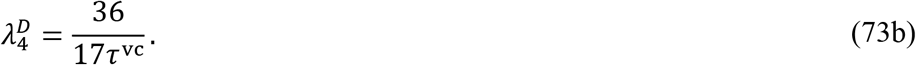

As Fig. 2 shows, the fusion dynamics is a stretched exponential function of time, occurring more slowly than the shape recovery dynamics of a droplet with a deformation represented by the lowest-order Legendre polynomial (“P2”). As the order of the Legendre polynomial increases, shape recovery becomes even faster. Hubstenberger et al. (Hubstenberger *et al.*, 2013) have observed that shape recovery of elongated grP-bodies was much faster than the fusion of grP-bodies. However, Fig. 2 shows that the results predicted for viscous droplets are unlikely to reach the two orders of magnitude difference in timescales, which Hubstenberger et al. suggested as indicating elasticity. Next we presents results for viscoelastic droplets.

**FIG. 2.**
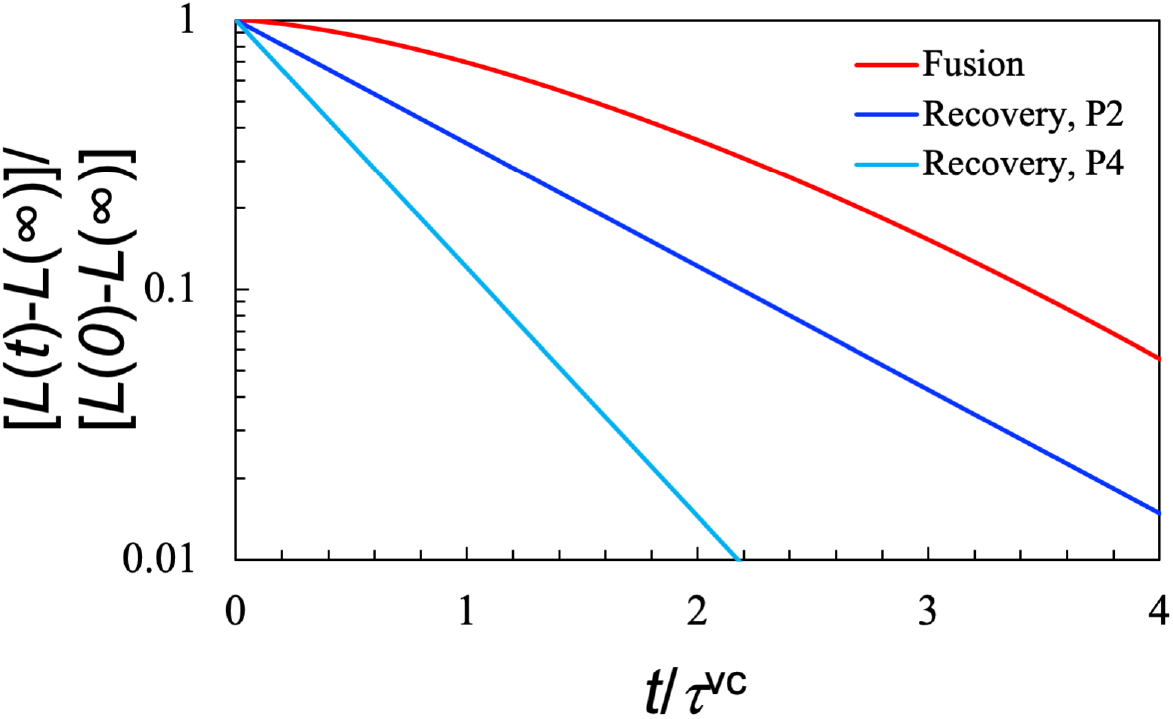
Comparison of droplet fusion dynamics and shape recovery dynamics. Fusion starts with two equal-sized droplets at contact and ends with a single larger spherical droplet having the same total volume. Deformation recovery is illustrated in Fig. 1; P2 and P4 indicate that the deformation is represented by the second- or fourth-order Legendre polynomial. *L* denotes the half-length of each system.

### Shape recovery of viscoelastic droplets

We now examine the time dependence of *f_l_*(*t*) for viscoelastic droplets in an ideal fluid. For the Maxwell model of linear viscoelasticity, substituting Eq. (59) into Eq. (63) yields

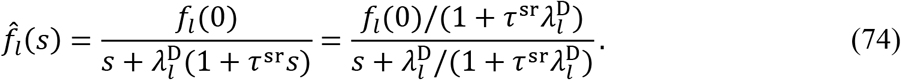

The inverse Laplace transform is

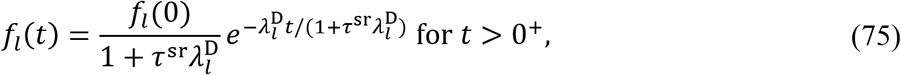

which is an exponential function of time. The recovery rate, 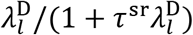, is less than the counterpart for a Newtonian fluid (when *τ*^sr^ = 0) by a factor of 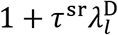. The apparent zero-time deformation, 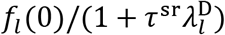, is also less than the nominal value, *f_l_*(0), by the same factor, as required by the conservation of area under the curve [see Eq. (66)]. So evidently there is a missing amplitude of 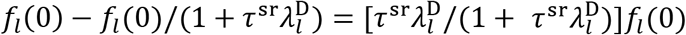; we elaborate its origin in Appendix B. This can be traced to *Ĝ*(∞) = 0 for the Maxwell model. The 0 value of *Ĝ*(∞) violates an assumption used for deriving Eq. (74), which is that *Ĝ*(*s*) is much greater than the exterior viscosity. The full solution without this assumption does not suffer from this artifact.

Next we consider the Jeffreys model. Substituting Eq. (60) into Eq. (63) gives

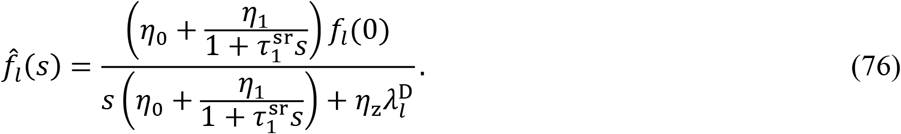

The inverse Laplace transform of Eq. (76) is a sum of two exponentials,

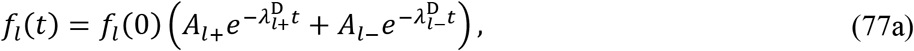

as opposed to a single exponential for a Newtonian or Maxwellian fluid. The recovery rates and corresponding amplitudes are

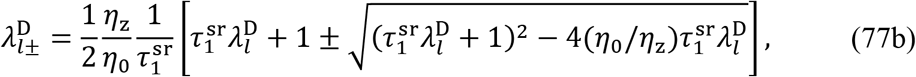

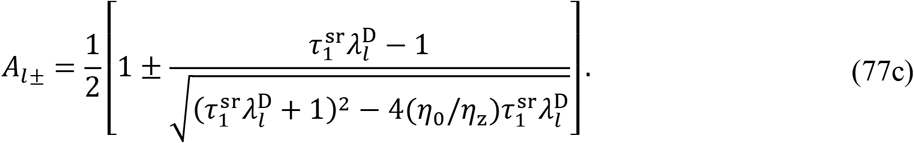

In the limit *η*_1_ → 0, the two recovery rates of Eqs. (77b) reduce to 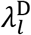 and 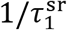, and the corresponding amplitudes given by Eq. (77c) reduce to 1 and 0, respectively. Thus Eq. (77a) properly reduces to the single exponential of Eq. (72) for a Newtonian fluid. In the limit *η*_0_ → 0, the two recovery rates become infinite and 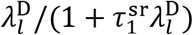, and the corresponding amplitudes are 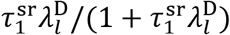 and 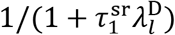. This result for the *η*_0_ → 0 limit thus reveals that the missing amplitude of the Maxwell model is due to an instantaneous decay at *t* = 0^+^, consistent with the analysis in Appendix B.

The scaled recovery rates 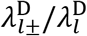 and the amplitudes *A*_*l*±_ depend only on two dimensionless parameters: *η*_0_/*η_z_* and 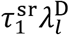. It can be easily shown that 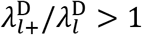 whereas 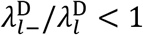 for all parameter values. For a given *η*_0_/*η_z_*, 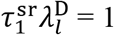 is a special point, where *A*_*l*+_/*A*_*l*−_ is 1 and 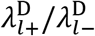 reaches its minimum. At this special 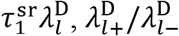 gets closer and closer to 1 as *η*_0_/*η_z_* approaches 1, i.e., when the Newtonian component of the complex shear modulus becomes dominant. In Fig. 3 we compare the shape recovery curves for *η*_0_/*η*_1_ = 5:1, 1:1, and 1:5 while holding 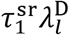 at 1. Whereas the curve at *η*_0_/*η*_1_ = 5:1 is close to the counterpart of Newtonian droplets, the curve at *η*_0_/*η*_1_ = 1:5 is clearly non-exponential, with a fast decay followed by a slow decay. The curve at *η*_0_/*η*_1_ = 1:1 is intermediate between those two.

**FIG. 3.**
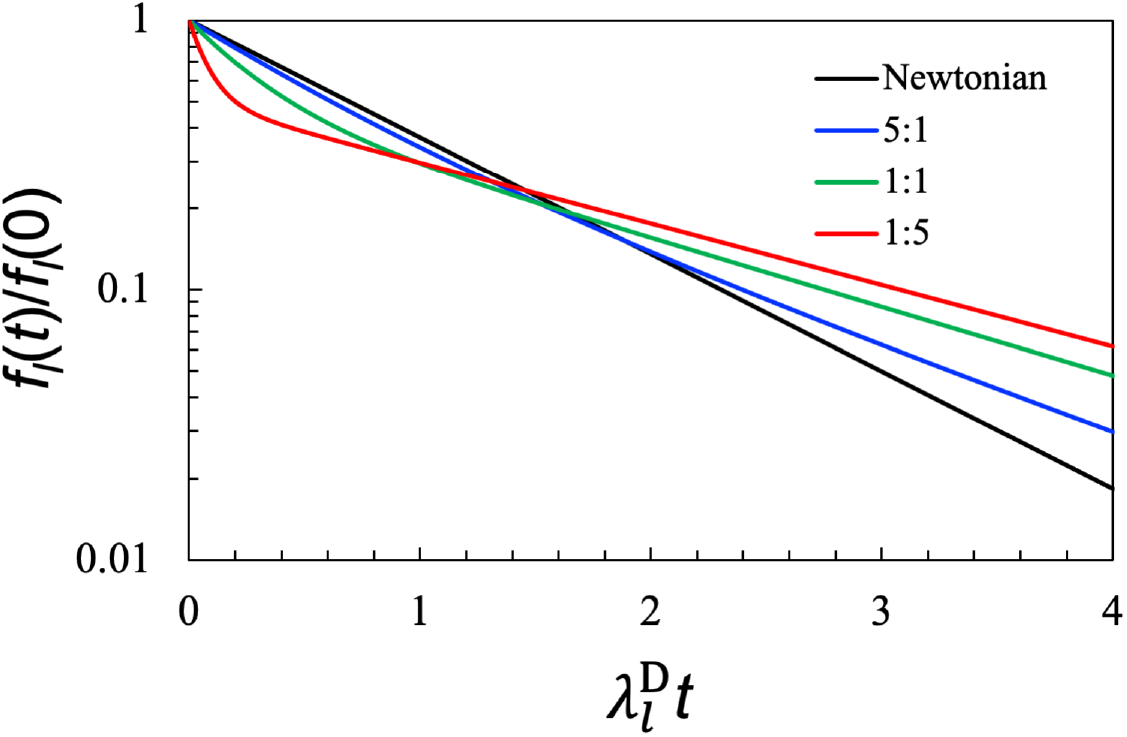
Effect of the *η*_0_/*η*_1_ ratio on the shape recovery dynamics of viscoelastic droplets in an ideal fluid. The shear relaxation rate is fixed at 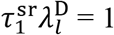, where the two decay components of *f_l_*(*t*) have equal amplitudes. The values of *η*_0_/*η*_1_ are shown in the legend. The result for a Newtonian droplet is displayed for reference. The results in the figure are valid for any *l*.

Recently we have found that modulating intermolecular interactions, e.g., by adding salt, can tune the *η*_0_/*η*_1_ ratio (Zhou, 2021). For PGL-3 protein droplets, the Maxwellian component dominates (i.e., *η*_0_/*η*_1_ < 1) at low salt while the Newtonian component dominates (i.e., *η*_0_/*η*_1_ > 1) at high salt. Qualitatively, we expect that shape recovery curves of PGL-3 droplets to become less non-exponential at increasing salt concentration. Experimental studies of salt effects on the shape recovery dynamics of biomolecular condensates will be of particular interest. Salts also have complex effects on the shear relaxation moduli of polymer blends (Akkaoui et al., 2020).

Finally let us consider the Burgers model. Substituting Eq. (61) into Eq. (63) yields

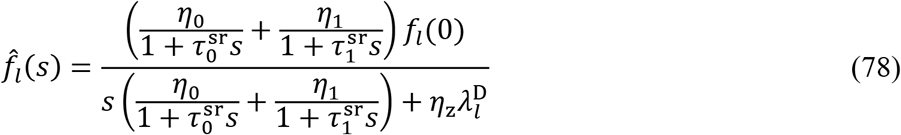

The inverse Laplace transform again gives *f_l_*(*t*) as a bi-exponential like Eq. (77a). The Burgers model has *Ĝ*(∞) = 0, and correspondingly there is also a missing amplitude, amounting to 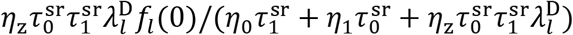. The two recovery rates and the corresponding amplitudes can be found in Appendix C.

The foregoing results for viscoelastic droplets in an ideal fluid also apply to the shape recovery dynamics of ideal-fluid bubbles in a viscoelastic medium, after replacing 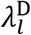 with 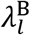. The expressions for these two relaxation rates are given in Eqs. (64) and (68), respectively. From here on, we will focus on the Jeffreys model of linear viscosity.

### Effect of shear relaxation rate

Let us look closely at the effect of 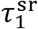, the shear relaxation rate in the Jeffreys model, on the shape recovery dynamics of viscoelastic droplets in an ideal fluid. In Fig. 4, we display shape recovery curves for 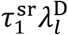 at 0.1, 1, 10, and 50 while holding *η*_0_/*η*_1_ at 1:5. Each *f_l_*(*t*) curve exhibits two successive decays. With increasing 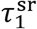, the amplitude of the first decay becomes larger and that of the second decay becomes smaller, while the recovery rates of both decays become slower. The latter trend is a sign for slowdown in shape recovery by shear relaxation.

**FIG. 4.**
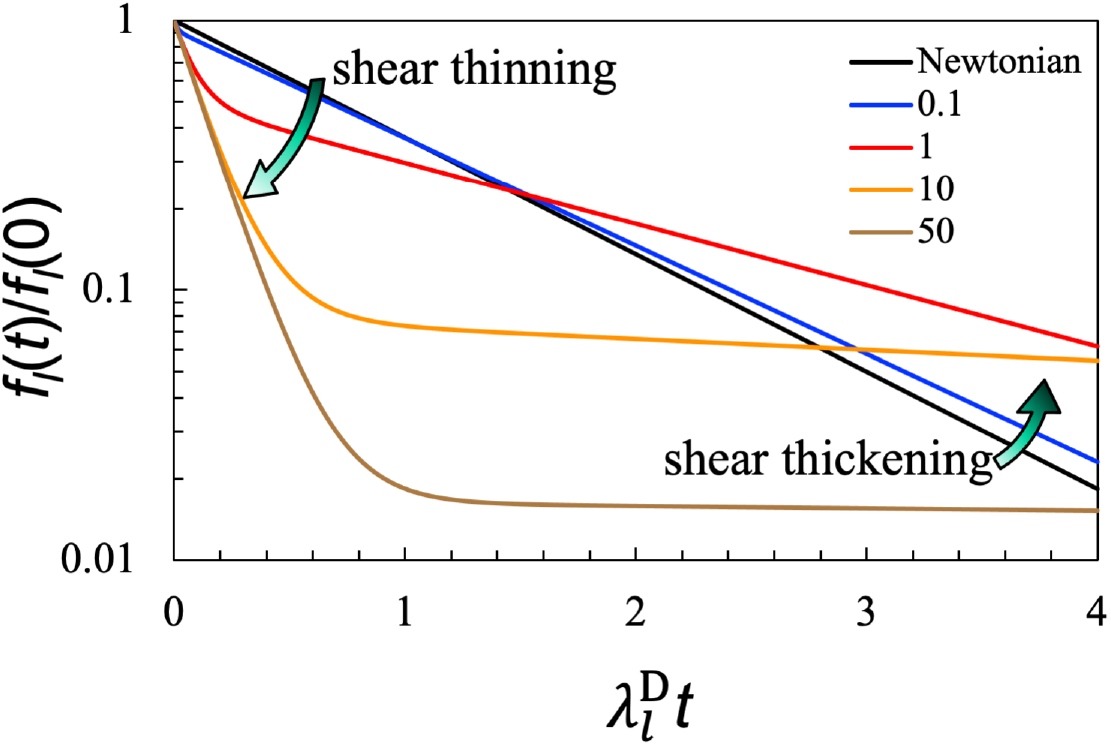
Effect of the shear relaxation rate on the shape recovery dynamics of viscoelastic droplets in an ideal fluid. The *η*_0_/*η*_1_ ratio is fixed at 1:5, whereas the values of 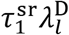 are shown in the legend. The result for a Newtonian droplet is displayed for reference. The results in the figure are valid for any *l*.

To understand these results more deeply, let us consider two extremes. For extremely fast shear relaxation, i.e., 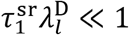, expanding Eqs. (77b) and (77c) in powers of 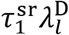 and keeping terms up to the first power, we find

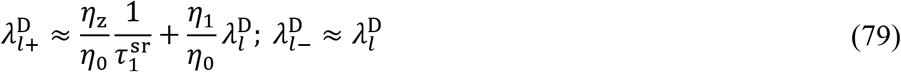

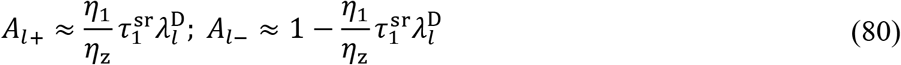

As illustrated by the curve at 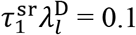 in Fig. 4, the first exponential, 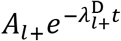, has a small amplitude and decays rapidly to 0, and hence is less important. The remaining exponential, 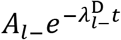, has an amplitude close to 1 and a recovery rate that is lower than but close to 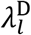, the Newtonian result. In other words, for 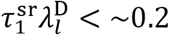, the shape recovery dynamics of viscoelastic droplets is almost a single exponential, but with a recovery rate slower than the Newtonian counterpart and thus apparently indicating shear thickening (i.e., increase in apparent viscosity).

In the opposite limit where shear relaxation is extremely slow, i.e., 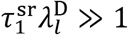, we expand Eqs. (77b) and (77c) in powers of 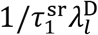 and keep terms up to the first power, finding

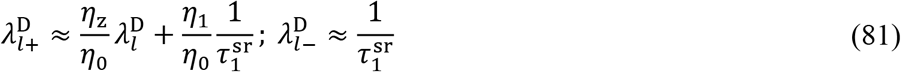

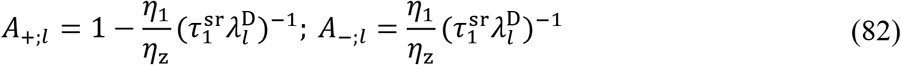

As illustrated by the curves at 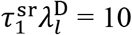 and 50 in Fig. 4, the first exponential has an amplitude that approaches 1 and a recovery rate that approaches 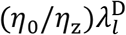, while the second exponential has a very small amplitude but a very slow recovery rate, 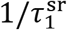. The ratio of the areas under the two exponentials is approximately *η*_0_/*η*_1_ and hence both areas can be significant, but the second, slow exponential may be difficult to detect experimentally due to the small amplitude. The apparent dominance of the first exponential thus gives the impression of fast shape recovery. Note that 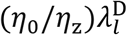 is the recovery rate of a Newtonian fluid with viscosity at *η*_0_. In essence, when shear relaxation is very slow, only the Newtonian component of the shear relaxation modulus is active in slowing down shape recovery (i.e., the Maxwell component of the shear relaxation modulus remains dormant on the timescale of the shape recovery). Therefore, during shape recovery, viscoelastic droplets appears to exhibit shear thinning (decrease in apparent viscosity) when shear relaxation is very slow. This apparent shear thinning could explain, at least partially, the shorter than expected timescale of grP-body shape recovery observed by Hubstenberger et al. (Hubstenberger *et al.*, 2013).

Shear relaxation could potentially present a new rate-limiting mechanism for shape recovery. One would then naively expect shape recovery to slow down with decreasing shear relaxation rate (i.e., increasing 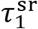). Indeed, as noted above, the two recovery rates 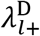 and 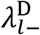 decrease with increasing 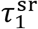. However, shear relaxation also affects the amplitudes of the two exponentials. Thus paradoxically the major exponential (i.e., the one with the larger amplitude) has a recovery rate lower than 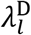 when shear relaxation is fast (e.g., 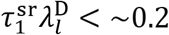) but a recovery rate higher than 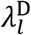 when shear relaxation is slow (e.g., 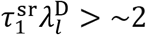) (Fig. 4). The opposite deviations from 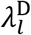 correspond to shear thickening and shear thinning, respectively. Interestingly, associative polymers, which may serve as a generic model of biomolecular condensates, exhibit shear thickening at moderate steady shear rates and shear thinning at high steady shear rates (Tripathi et al., 2006). We propose that the shear thickening and thinning phenomena of viscoelastic droplets during shape recovery and of associative polymers under steady shear have a common explanation. Our recent experimental studies that dissected droplet fusion data have presented evidence for shear thickening and shear thinning of biomolecular condensates (Ghosh *et al.*, 2021; Ghosh and Zhou, 2020).

Of course we should not forget that there is also a minor exponential, which starting at 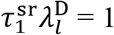 has a time constant that becomes dominated by shear relaxation and less and less dependent on interfacial tension. For extremely slow shear relaxation, the time constant becomes 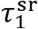 itself, thus directly showing that shear relaxation rate-limits shape recovery. However, this new mechanism only accounts for a very small fraction of the total amplitude, thus making experimental verification challenging.

As noted in the preceding subsection, the results illustrated by Fig. 4 are also applicable to the shape recovery of bubbles in a viscoelastic medium upon replacing 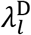 with 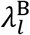. Ali and Prabhu (Ali and Prabhu, 2019) recently presented such data where the viscoelastic medium is coacervates formed by potassium poly(styrenesulfonate) and poly(diallyl dimethylammonium bromide). They observed an initial decay followed by a slow second decay, qualitatively in agreement with our prediction. The shear relaxation in their system has 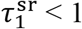 s and becomes faster with increasing salt (similar to results found for PGL-3 droplets (Zhou, 2021)). In comparison, the time constants of their second decay, corresponding to our 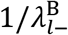, are mostly > 1 s. Ali and Prabhu analyzed their data essentially by assuming 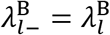, i.e., completely neglecting shear relaxation. However, even fast shear relaxation can affect 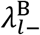, and hence its neglect can lead to underestimation of the interfacial tension. For example, at 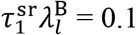, we predict 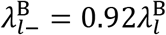. Ali and Prabhu also reported the time, *t*_e_, for the initial decay to complete. *t*_e_ should scale with 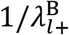, i.e., the time constant of the initial decay, which we predict to be significantly affected by the shear relaxation rate [see 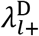 given by Eq. (79)]. Indeed, Ali and Prabhu reported an approximately 10-fold decrease in *t*_e_ when the salt concentration increased from 1.55 to 1.85 M. Over the same range of salt concentration, the time constants of their decay changed by < 2-fold whereas the shear relaxation rates changed substantially, suggesting their major role in determining *t*_e_.

### Finite viscosity ratio: Newtonian fluids

So far we have only considered cases where either the exterior or the interior fluid is modeled as ideal. Now we want to present cases where the viscosity ratio between the two phases is finite so that both interior and the exterior fluid dynamics must be treated at the same time. As a prelude to cases where a viscoelastic fluid interfaces with a Newtonian fluid, here we first consider the case where both phases are Newtonian fluids. The shape recovery dynamics is an exponential function [see Eq. (53)], with the recovery rate *λ_l_* given by Eq. (20a). For *l* = 2, the recovery rate is

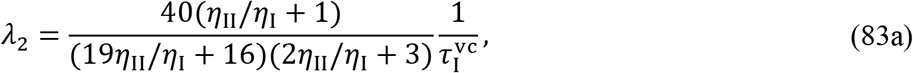

where

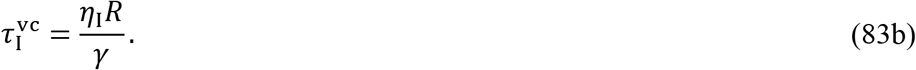

Equation (83a) is identical to the recovery rate widely quoted for droplets approximated as ellipsoids (Ali and Prabhu, 2019; Luciani *et al.*, 1997; Minale, 2010; Mukherjee and Sarkar, 2010; Yu *et al.*, 2004).

Although shapes specified by the second-order Legendre polynomial (“P2”) have been assumed to be equivalent to ellipsoids (e.g., Benayad et al., 2021), there are actually noticeable differences between them. As detailed in Appendix D, P2 combined with a small amplitude fourth-order Legendre polynomial (P4) does an excellent job in reproducing ellipsoidal shapes. A deformation with an ellipsoidal shape thus requires a linear combination of *f*_2_(*t*) and *f*_4_(*t*) (see Fig. 2), leadging to a bi-exponential decay even when both phases are Newtonian fluids. For the rest of the paper, we will limit to *l* = 2, focusing attention instead on non-exponentiality arising from viscoelasticity.

In Fig. 5(a) we present the shape recovery curves for Newtonian droplets in a Newtonian fluid with *η*_II_/*η*_I_ at 10, 1.5, and 0.5. This will serve as references for assessing the effect of droplet viscosity in the next subsection. One can tune *η*_II_/*η*_I_ by changing temperature or salt concentration. In cases where the dense phase is inside droplets, *η*_II_/*η*_I_ is much higher than 1 while away from the critical point, and approaches 1 as the critical point is reached. For complex biomolecular condensates, one can also tune *η*_II_/*η*_I_ by varying their macromolecular composition (Elbaum-Garfinkle *et al.*, 2015; Ghosh and Zhou, 2020; Zhang *et al.*, 2015).

**FIG. 5.**
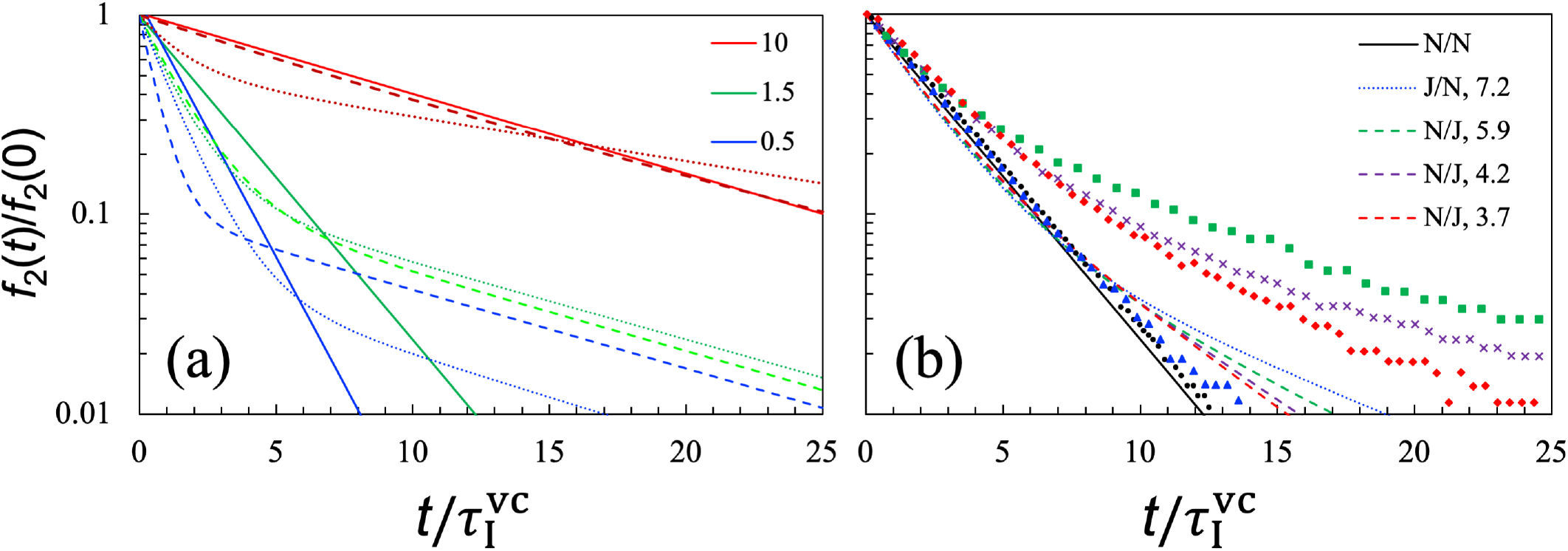
Shape recovery curves at a finite viscosity ratio between the droplet and bulk phases. (a) Comparison of *f*_2_(*t*) for three cases of pairing interior and exterior fluids: Newtonian with Newtonian (solid curves), viscoelastic with Newtonian (dotted curves), and Newtonian with viscoelastic (dashed curves). The viscosity ratios, i.e. *η*_II_/*η*_I_, are shown in the legend. Viscoelasticity is given by the Jeffreys model, with *η*_0_/*η*_1_ = 1:5 and 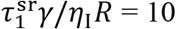. (b) Comparison of our analytical results (curves) with Verhulst et al.’s experimental data (symbols) for five systems, all with *η*_II_/*η*_I_ = 1.5. The viscoelasticity was reported as fitting to the Oldroyd B model, with *η*_0_/*η*_1_ ≈ 2:1 and 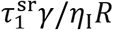 indicated in the legend. Curves and symbols have matching colors for the same systems.

### Finite viscosity ratio: viscoelastic fluids

To determine the shape recovery dynamics of viscoelastic droplets in a Newtonian fluid, we obtain *f_l_*(*t*) of Eq. (69a) by numerical Laplace inversion (Stehfest, 1970). The results are displayed as dotted curves in Fig. 5(a) for Jeffreys droplets with *η*_II0_/*η*_II1_ = 1:5, 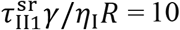, and *η*_II_/*η*_I_ = 10, 1.5, and 0.5. Similar to the results presented in Figs. 3 and 4 for *η*_II_/*η*_I_ = ∞, for each finite *η*_II_/*η*_I_ value, the initial decay of *f_l_*(*t*) for the Jeffreys droplet (“J/N”) is faster than that of the corresponding Newtonian droplet (“N/N”). The J/N curve then slows down and crosses the N/N curve.

It is interesting to compare the shape recovery curves between Jeffreys droplets in a Newtonian fluid (i.e., “J/N”) and Newtonian droplets in a viscoelastic medium (“N/J”). The N/J results are shown as dashed curves in Fig. 5(a). When *η*_II_/*η*_I_ > 1, relative to the J/N curves, the corresponding N/J curves move closer to the N/N curves. Conversely, when *η*_II_/*η*_I_ < 1, it is the J/N curves that are closer to the N/N curves. At *η*_II_/*η*_I_ = 1.5 and 0.5, with 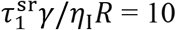, the shear relaxation of the viscoelastic fluids occurs on a timescale that is apprximately 10 times longer than both the interior and exteruor viscocapillary times. Correspondingly, the shape recovery curves at *η*_II_/*η*_I_ = 1.5 and 0.5 all have a slow decay with a time constant close to 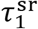. So once again we observe the scenario where the slow decay is dictated by shear relaxation and independent of interfacial tension.

In Fig. 5(b), we compare our analytical solution with the experimental data of Verhulst et al. (Verhulst *et al.*, 2009a) for N/N, J/N, and N/J systems, all with *η*_II_/*η*_I_ = 1.5. For the N/N system, there is good agreement between theory and experiment. For the J/N system, the analytical curve has a significant late slow decay whereas the experimental curve deviates only slightly from the counterpart for the N/N system. For the N/J systems, both the experimental and analytical curves have a long-time tail and the decay rate of the tail decreases with increasing 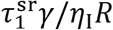, indicating that this part of the curves is dominated by shear relaxation. However, the analytical solution significantly underestimates the amplitude of the tail. We attribute the underestimation to the fluid-dynamics model adopted here for Newtonian droplets inside a viscoelastic fluid, not our analytical solution. The experimental N/J curves are above the corresponding N/N curve at all times, but the analytical solution of the present fluid-dynamics model predicts that the N/J curves must intersect the N/N curve and all the curves integrate to the same total area. Clearly the fluid-dynamics model needs to be modified for the N/J systems in order to achieve quantitative agreement with the experimental data. While phenomenological models have achieved partial success in fitting the experimental data (Minale, 2010; Yu *et al.*, 2004), we believe that it is more important to identify physical ingredients missing from the fluid-dynamics model and then solve the physical model exactly.

We also compared our analytical solution with the numerical solution of Hooper at al. (Hooper *et al.*, 2001), who used the Oldroyd B model for viscoelasticity. The analytical and numerical results agree well for Newtonian droplets in a Newtonian fluid. However, for viscoelastic droplets in a Newtonian fluid, Hooper et al.’s initial decay of the shape deformation is faster. Although the analytical solution is limited to linear order (specifically in the extent of shape deformation and in the linearization of the Oldroyd B model), we suspect the discrepancy is largely due to errors in the numerical solution. The Oldroyd B model is known to be difficult to implement numerically and to be prone to generate numerical instability (Verhulst *et al.*, 2009b). Our own recent numerical solution using the COMSOL software has shown close agreement with the analytical solution (Naderi, Peng & Zhou, to be published). The analytical solution thus presents a unique benchmark for testing the accuracy of numerical solutions.

## CONCLUDING REMARKS

We have presented an exact analytical solution for the recovery dynamics of biomolecular droplets from small-amplitude deformation. Whereas viscocapillarity sets the timescale for the shape recovery dynamics of viscous droplets, with viscoelasticity, shape recovery becomes multi-exponential. For the Jeffreys model of viscoelasticity featuring a single shear relaxation rate, two exponentials are predicted for the shape recovery dynamics, with time constants shorter and longer, respectively, than the viscocapillary timescale. Shear relaxation inside the droplets affects both the time constants and amplitudes of the exponentials, with one exponential becoming dominant under some conditions. For moderately fast shear relaxation, the dominant exponential is the one with a time constant longer than the viscocapillary timescale, which can be interpreted as an apparent increase in viscosity. Conversely, for very slow shear relaxation, the dominant exponential is the one with a time constant shorter than the viscocapillary timescale, which can be interpreted as an apparent decrease in viscosity. Therefore, during shape recovery, viscoelastic droplets exhibit shear thickening at fast shear relaxation rates but shear thinning at slow shear relaxation rates. Under the latter condition, the time constant of the minor exponential is dictated by shear relaxation, which can thus be seen as a new rate-limiting mechanism for shape dynamics.

We anticipate qualitatively similar effects of shear relaxation on droplet fusion dynamics. Indeed, preliminary data from COMSOL calculations on droplet fusion and shape recovery support this contention (Naderi, Peng & Zhou, to be published). Therefore, in droplet fusion, we can also expect shear thickening for condensates with fast shear relaxation rates but shear thinning for condensates with slow shear relaxation rates. This expectation is confirmed by our recent experimental studies that dissected droplet fusion data of biomolecular condensates (Ghosh *et al.*, 2021; Ghosh and Zhou, 2020).

There is great interest in condensate aging, especially in its connection with neurodegeneration, but there is little physical understanding of this phenomenon. One possible physical change of biomolecular condensates over time is the slowdown in shear relaxation (Jawerth *et al.*, 2020). Our analytical solution predicts that a slowdown in shear relaxation can lead to a very slow decay in shape dynamics, thereby giving the appearance of stalled or incomplete shape recovery and fusion. The work here thus reveals shear relaxation as a crucial link in understanding condensate aging.

Our analytical solution should prove useful for validating numerical solutions of fluid-dynamics equations for condensate shape changes. Such validation is important, as numerical solutions may be the only option for the theoretical treatment of nonlinear issues such as large-amplitude deformation and nonlinear constitutive relations (e.g., the Oldroyd B model). We do expect that our present solution is still qualitatively and even semi-quantitatively correct when nonlinearity is present, and as such will be good for fitting experimental data on shape recovery. Finally shape recovery and fusion may potentially be modeled by molecular simulations (Benayad *et al.*, 2021) and our analytical theory can motivate and guide data analysis in such studies.

## ACKNOWLEDGMENTS

This work was supported by Grant GM118091 from the National Institutes of Health.

## APPENDIX A: SOLUTION WHEN BOTH *η*_I_ AND *η*_II_ ARE FINITE

When both *η*_I_ and *η*_II_ are finite, we need to consider both the interior and exterior fluid dynamics. The boundary conditions on the interface between the droplet and bulk phases consist of the continuity of the velocity field,

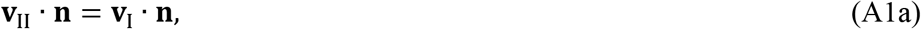

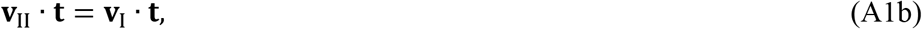

the kinematic equation,

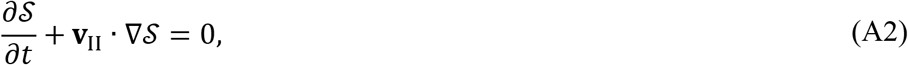

and the force-balance equations,

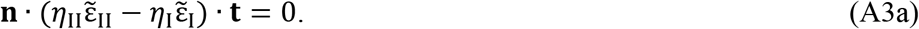

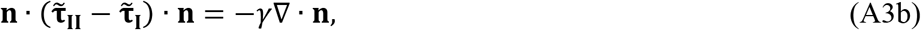

The interior velocity and pressure fields are given by Eqs. (31) – (33); the exterior counterparts are given by Eqs. (44) – (46), except that the term –*γ/ηR* on the right-hand side of Eq. (46) should be removed, since the effect of this term is already accounted in Eq. (33) for the interior pressure. Equations (A1a) and (A1b) lead to

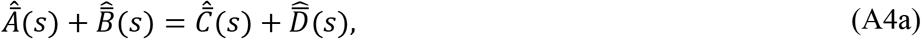

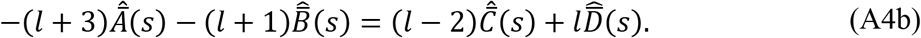

Equation (A3a) leads to

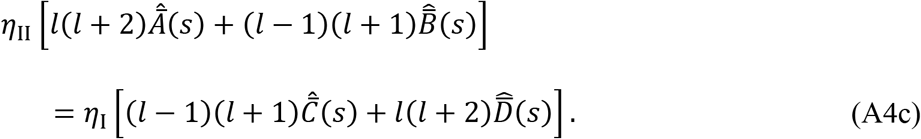

Equation (A3b) leads to

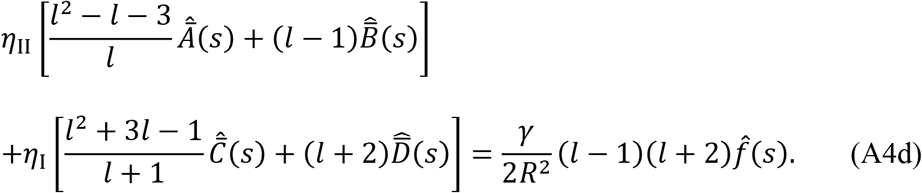

Finally Eq. (A2) leads to Eq. (41b), which is

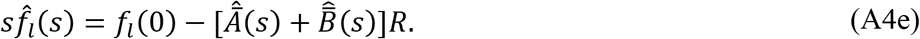

Solving Eqs. (A4a–e), we find the solution for 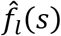 given Eq. (53), with the recovery rate *λ_l_* given by Eq. (20a).

## APPENDIX B: MISSING AMPLITUDE OF MAXWELL DROPLETS

To find the origin of the missing amplitude of Maxwell droplets inside an ideal fluid, let us look at 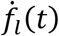, the rate of change in *f_l_*(*t*). Its Laplace transform is [using Eq. (41b) and Eqs. (42a, b) with *η* replaced by *Ĝ*(*s*)]

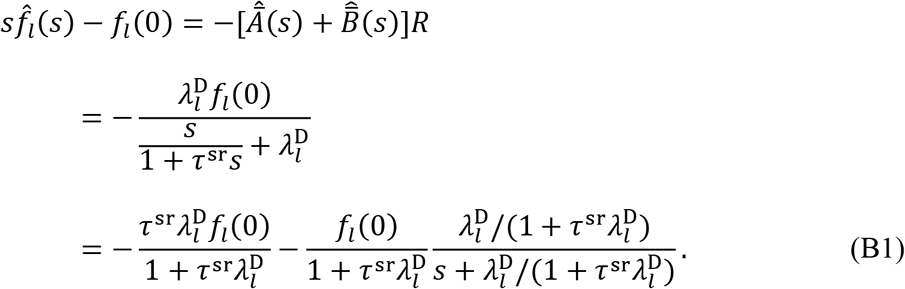

The inverse Laplace transform is

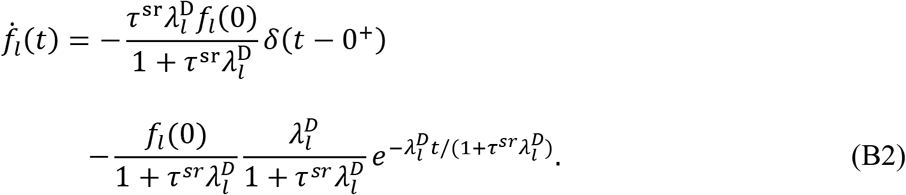

Upon integrating Eq. (B2) over time, the first term, a delta function, leads to an instantaneous drop in the amplitude at *t* = 0^+^, which is exactly the missing amplitude; the second term yields the result in Eq. (75), which is applicable for *t* > 0^+^.

In general, the behavior of *f_l_*(*t*) near *t* = 0 is dictated by 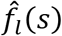 at *s* → ∞. If *Ĝ*(∞) is a nonzero finite value, then the asymptote of 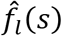 given by Eq. (63) is *f_l_*(0)/*s*, which upon inverse Laplace transform yields the correct value, *f_l_*(0), for *f_l_*(*t*) at *t* = 0. However, this analysis breaks down when if *Ĝ*(∞) = 0, as is the case for the Maxwell and Burgers models, leading to the missing amplitude. Note that the 0 value of *Ĝ*(∞) violates an assumption used for deriving Eq. (63), which is that *Ĝ*(*s*) is much greater than the exterior viscosity. The full solution, given by Eq. (69a), that accounts for the exterior viscosity does not suffer a missing amplitude. When *ĝ*_II_(∞) = 0 as in the Maxwell and Burgers models, Eq. (69c) still yields a finite *h_l_*(∞) and therefore Eq. (69a) gives the correct asymptote *f_l_*(0)/*s* for 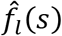.

## APPENDIX C: SHAPE RECOVERY DYNAMICS OF BURGERS DROPLETS

For viscoelastic droplets modeled by the Burgers model, the inverse Laplace transform of Eq. (78) yields the bi-exponential form of Eq. (77a) for the shape recovery dynamics. The two recovery rates and the corresponding amplitudes are

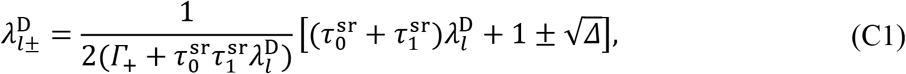

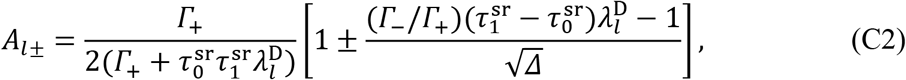

where

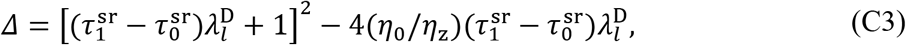

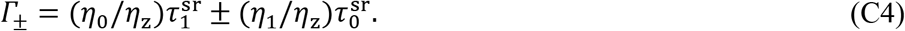

Note that

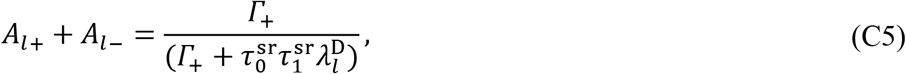

which is less than 1. So here also there is a missing amplitude, due to the fact that *Ĝ*(∞) = 0 for the Burgers model.

## APPENDIX D: EFFECT OF INITIAL DEFORMED SHAPE

Both Fig. 2 and Eqs. (73a, b) make it clear that the precise shape of the initial deformation affects the recovery dynamics. In particular, an ellipsoid can be approximated well by a combination of the second- and fourth-order Legendre polynomials, as illustrated in Fig. 6(a) for an ellipsoid with a half-length 1.4Æ and a half-width of 0.87*R*. In this case the P2 and P4 amplitudes determined by representing the elliptical cross section by a sum of Legendre polynomials are 0.305 and 0.073, respectively. Recovery dynamics from such an initial ellipsoidal shape can be described by a corresponding linear combination of *f*_2_(*t*) and *f*_4_(*t*), leading to a bi-exponential decay [Fig. 6(b)].

**FIG. 6.**
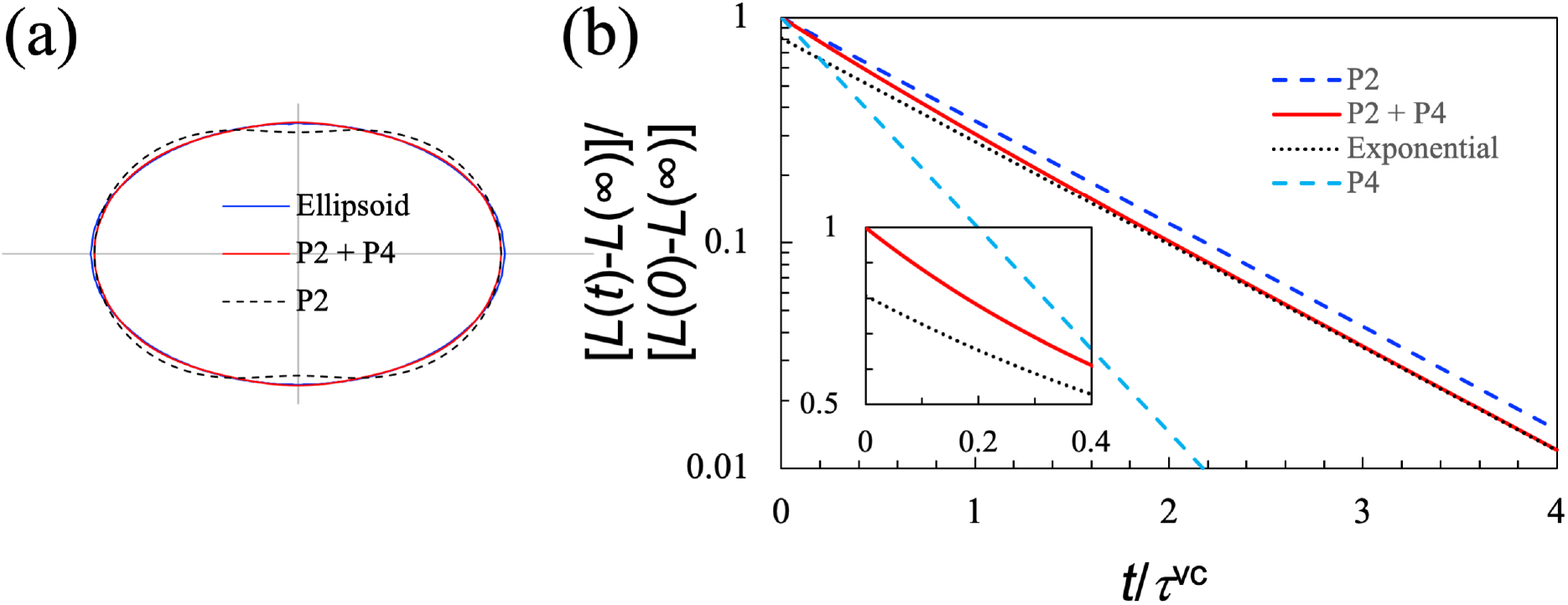
Effect of initial shape on recovery dynamics. (a) Representation of an ellipse (with major and minor semi-axes at 1.4 and 0.87) by the sum of the first, second (P2), and fourth-order (P4) Legendre polynomials. The amplitudes of P2 and P4 are 0.305 and 0.073, respectively. (b) Comparison of the deformation decay curves for P2, P4, and their combination (“P2 + P4”) that represents the ellipse in panel (a). The dotted curve labelled “Exponential” displays the P2 component in the P2 + P4 curve. Inset: difference between the P2 + P4 curve and the P2 component at short times. The ordinate is in linear scale, in contrast to the log scale in the main figure.

